# Complement active human and porcine serum induces natural competence for genetic transformation in the emerging zoonotic pathogen *Streptococcus suis*

**DOI:** 10.1101/2020.03.19.996926

**Authors:** Maria Laura Ferrando, Alex Gussak, Saskia Mentink, Peter van Baarlen, Jerry Mark Wells

**Affiliations:** Host-Microbe Interactomics, Animal Sciences, Wageningen University, Wageningen, The Netherlands

## Abstract

The acquisition of novel genetic traits by natural competence is a strategy used by bacteria in microbe-rich environments including animal or human hosts where microbial competition, antibiotics and host immune defences threaten their survival. We show here that several virulent strains of *Streptococcus suis*, an important porcine pathogen and zoonotic agent, become naturally competent for genetic transformation with plasmid or genomic DNA when cultured in active porcine and human serum, but not when it is pre-heated for 30 minutes at 56°C to inactivate complement. Competence is also not induced in active fetal bovine serum, which contains less complement factors and immunoglobulins than adult serum. Late competence genes, encoding the uptake machinery for environmental DNA, were highly upregulated in active serum. Competence development was independent of the early competence regulatory switch suggesting the presence of an alternative stress-induced pathway for regulation of the transformasome, a type 4-like pilus DNA binding and transport apparatus.

## Introduction

Natural competence, the process by which bacteria acquire environmental DNA and integrate into their genome [1] may confer selective advantages to bacterial pathogens during infections and lead to the emergence of hypervirulent pathogens [2]. In some streptococcal species, competence can be induced in laboratory medium by addition of a peptide pheromone [3] and in the environment by stress conditions as DNA-damage, nutrient limitation and antibiotics [4]. Factors inducing natural competence during infections are largely unknown.

*Streptococcus suis* (SS), a swine pathogen and emerging zoonotic agent which causes meningitis and septicemia [5], invades host tissues and spreads systemically through the bloodstream. Serotype 2 (SS2) includes highly pathogenic and zoonotic strains whereas SS14 embraces potentially zoonotic strains [5]. SS1, SS*7* and SS9 serotypes include strains highly virulent in pigs [6]. The high genomic diversity of SS suggests high levels of recombination between strains and is considered a risk factor for the emergence of novel hypervirulent and zoonotic strains [7]. We investigated the induction of natural competence in different serotypes of virulent SS strains under conditions that mimic infection-associated with serum factors during tissue invasion.

Active (not heat-treated) Porcine Serum (aPS) from blood isolated from sows contains innate and adaptive immune defense factors including the complement system, antimicrobial peptides and specific anti-SS antibodies [8]. We hypothesized that the general stress response of SS exposed to serum factors may induce natural competence [4]. Extracellular pheromone peptide, XIP (*com<u>X</u>*-<u>I</u>nducing <u>P</u>eptide) induces competence in SS [9]. XIP activates the ComR regulator that transcribes the master competence regulator *comX*, which in turn induces expression of the competence machinery (transformasome) including a type IV-like pilus [9]. Not all SS strains can be transformed by their cognate XIP including virulent SS*7* isolates [9], thus other methods for inducing natural competence would open up possibilities for efficient genetic manipulation.

## Materials and Methods

### Bacterial strains, plasmids and culture conditions

Bacterial strains and plasmids are listed in Table 1. SS strains were grown in Todd Hewitt Broth (THB; Oxoid) supplemented with 0.2% Yeast extract (Difco) (THY) at 37 ⁰C with 5% CO_2_. Porcine serum (PS) (cat. <u>26250084</u>, Gibco), Fetal Bovine Serum (FBS) (cat. FBS-12A, Capricorn) was slowly defrosted at 4 ⁰C overnight (O/N), split in aliquots of 30 ml and stored at −20 ⁰C for extended use. Human Serum (HS) was derived from blood that was drawn from three healthy volunteers and collected in X tubes. Blood was allowed to clot for 15 min at room temperature, serum was collected after centrifugation for 10 min at 4000 g at 4°C, pooled and subsequently stored at −20 ⁰C. Heat-inactivated serum was prepared by incubation of serum for 30 min at 56 °C.

**Table 1:**
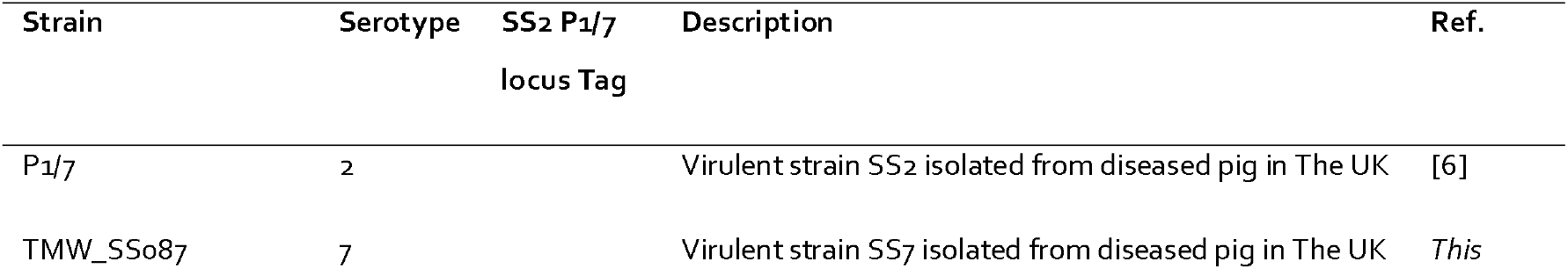
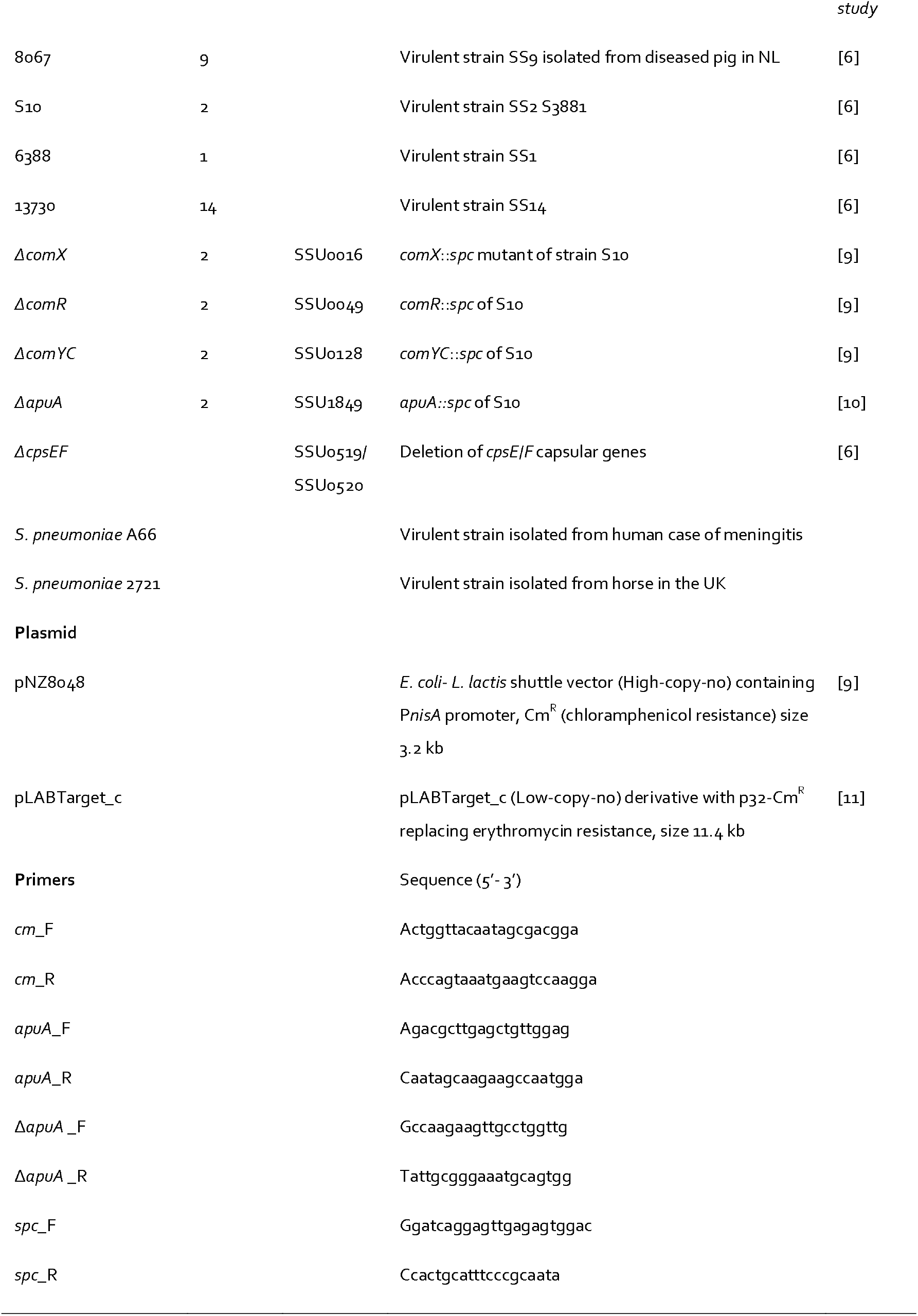
strains and plasmids used in this study.

When necessary, antibiotics were added to culture media at the following concentrations: chloramphenicol (Cm) 5 μg/ mL and spectinomycin (Spc) 100 μg/ mL.

### RNA-seq

Bacterial cultures were grown in triplicate (10ml each) to the mid-exponential phase in either THY medium or serum. After centrifugation, the bacterial pellet was resuspended in QIAzol buffer (Qiagen), added to vials containing silica beads and lysed using FastPrep-24 5G (MP Biomedicals) (settings: 4.0 m/sec and run time 40 seconds). The supernatant obtained by 5 minutes of centrifugation at 16.000 g of bacterial lysate was stored at −80 °C or immediately purified with the miRNeasy mini kit (Qiagen). RNA quantity was determined using the Qubit 4 Fluorometer and Nanodrop spectrophotometer (Thermofisher), while RNA quality expressed in RIN (RNA Integrity Number) was determined by electrophoresis using the 2200 TapeStation system (Agilent). Illumina mRNA sequencing (RNA-seq) was carried out at Novogene Bioinformatics Technology Co., Ltd., in Hong Kong, China. Strand-specific libraries were prepared and sequencing by Hiseq Illumina PE150 platform was used to generate short reads. FASTQ reads were analysed using the CLC Genomics Workbench 11.0 (Qiagen). After trimming and quality check, reads of both strains were mapped against the SS2 P1/*7* reference genome (<u>AM946016</u>). The expression levels were normalized as Transcripts Per Million (TPM) using the RNA-seq analysis option in CLC GWB. For the identification of differentially expressed transcripts we used a two-tailed t-test with adjusted p-values using false discovery rate (FDR). Genes with adjusted p-values below 0.05 were considered statistically significantly differentially expressed. Gene expression was expressed as log_2_.

### Natural competence acquisition assay

3 ml O/N bacterial culture in THY were used to inoculate 27 ml of serum (10-fold dilution, OD600~0.08). The inoculated serum was incubated at 37 ⁰C for 4 hours. Every hour, 200 μl of bacterial subculture in serum was transferred into 1.5 ml Eppendorf tubes and 10 μl of plasmid (pNZ8048 or pLABTarget_c) with a concentration of 200 ng/μl was added to the subculture that was successively incubated at 37 ⁰C for two additional hours to permit the acquisition of plasmid. After that, the subculture was plated on THY agar plates with the required antibiotic and incubated at 37 ⁰C for 1 or 2 days to permit the outgrowth of possible recombinant colonies. In most of the experiments, a possible competence acquisition was also verified in laboratory media growing SS strains in THY culture and subculture following the same experimental conditions as previously described. Furthermore, to verify that bacteria had not previously acquired natural antibiotic resistance, 10 μl of sterile H_2_O was added to the bacterial subculture as negative control. A WT strain was tested as control when Δ*com* mutants were assed for competence. Putative transformed colonies were verified by colony PCR employing *cm*_F/*cm*_R primers (Table 1) to validate acquisition of the corresponding plasmid carrying the Cm antibiotic resistance marker.

To verify whether a homologous recombination event had occurred in aPS after co-incubation of SS and plasmids, a PCR fragment (primers: *apuA*_F/*apuA*_R) obtained from a Δ*apuA* mutant [10] was added to 200 μl of 10-fold O/N subculture in aPS as described above. Bacterial lysate of this mutant was derived from 30 ml of O/N culture was obtained, concentrated in 2 PBS ml after centrifugation, heated it at 99 ⁰C for 15 min, and then further clarified at high centrifugation speed (16.000 g for 5 min). Clarified bacterial lysates were plated on THY agar and incubated at 37 ⁰C to verify the absence of viable cells. 20, 50 and 100 μl of clarified bacterial lysates were used in transformation assays using SS cultured in aPS as previously described. Correct homologous recombination in SS2 P1/*7* genome, in which the *apuA* gene is interrupted by a spc cassette, was verified by colony PCR using the primers Δ*apuA* _F/Δ*apuA* _R and *spc*_F/*spc*_R (Table 1).

The low Molecular Weight (MW) fraction containing short peptides that might include XIP, was separated by centrifugal ultrafiltration of 20 ml aPS using a filter with 10 kDa cut-off (Vivaspin^®^ 20) at 8000 *g* for 1 hour, and 10 ml of filtrated low-MW serum fraction where used in the transformation assays.

To test whether the availability of iron could influence the transformation efficiency in serum, an O/N culture of SS was first incubated at 37 ⁰C in THY with the addition of 100 μM deferoxamine mesylate salt (DIF) [12] (D9533-1G, Sigma) and inoculated the day after in 30 ml aPS containing DIF at the same concentration. To test an iron-enriched condition 100 μM Iron (Fe^2+^) sulfate heptahydrate [12] (Sigma) was added to aPS.

The induction of competence under restricted nutrient conditions were tested was assayed in Complex Media (CM) without the addition of glucose as previously described [10].

In every experiment, bacterial growth in serum was measured at OD_600_ (SpectraMax M5); every hour the serum culture was diluted 1:1 in PBS.

## Results and discussion

SS growth was observed in all sera (Figure 1, Table S1). Transcriptomics of reference virulent SS strains (i.e. SS2 strain P1/*7* and SS9 strain 8067) grown in aPS revealed extensive differential gene expression compared to THY medium. Transcriptome data are available via https://doi.org/10.17026/dans-z5w-v5m3. For SS2, most of the late competence DNA uptake machinery genes were upregulated in serum (log2-fold-change >2.0, Table 2). In both strains *comYC* encoding the major DNA-binding pilus protein was the most upregulated gene in serum (log2-fold-change 5.48 for SS2 and 4.36 for SS9). The genes *comR* and *comX*, regulators of competence induction and homologous recombination were downregulated in aPS compared to THY (Table 2). A predicted fratricide protein (SSU1911) and a bacteriocin gene cluster (SSU0039-SSU0045) [13] were also expressed in aPS (log2-fold-change between 0.7 and 2.0) suggesting that non-competent bacteria might be lysed [13], [14], [15], [12].

**Figure 1.**
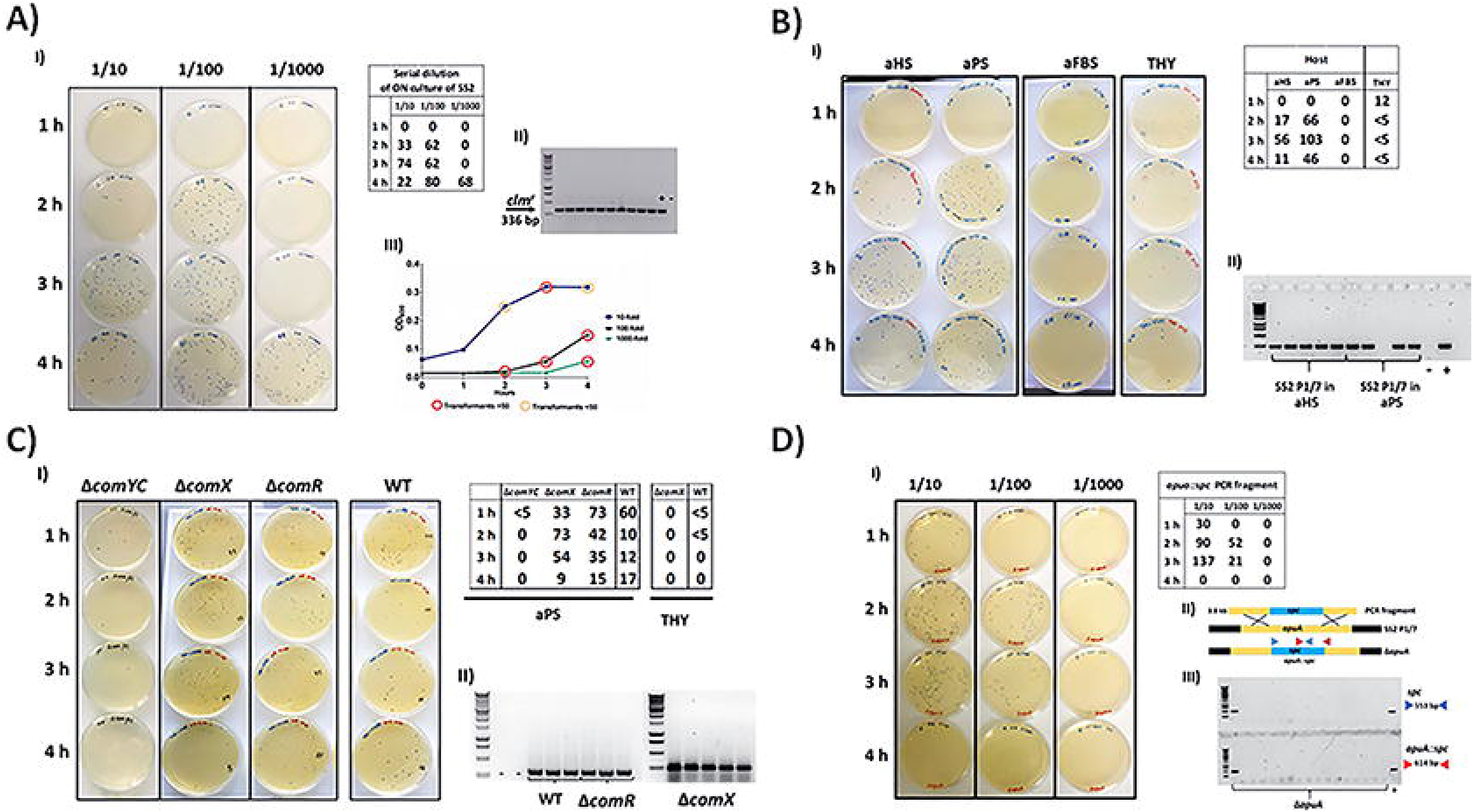
Induction of natural competence in *S. suis* during growth in serum. A) Induction of natural competence of serial 10-fold dilution of SS2 P1/*7* O/N culture inoculated in 30 ml of aPS: In the table are reported the number of transformants recovered by plating 200 μl of subculture of serum on selective THY plates; II) PCR detection of plasmid uptake by competent SS2 P1/*7*. The presence of a band at 336 bp indicates correct transformants positive for the Cm^r^ marker; III) bacterial growth curves combined with number of transformants. B) Natural competence acquisition tested in aPS, aHS, FBS and THY and scoring of the putative transformants of each growth condition reported in the table; II) a colony PCR to verify the uptake of exogenous plasmid DNA (336 bp) by SS2 P1/*7*. C) Transformation efficiencies of the master regulators ComR, ComX and transposome ComYC mutants in SS2 strain S10 grown in aPS and THY (THY plates not shown). II) colony PCR to verify the uptake of exogenous plasmid DNA (336 bp) by SS2 S10 WT and SS2 S10 Δ*comR*/Δ*comX* mutant strains. D) Transformation of SS2 P1/*7* with gDNA from Δ*apuA* mutant strain^12^. The table number reported the number of candidate transformants recovered on THY agar containing spectinomycin. II) representation of the homologous recombination event using a 3.8 kb PCR fragment from the genomic DNA of SS2 P1/*7* Δ*apuA*^12^ with the *apuA* gene fragment of WT SS2 P1/*7* (2.8 kb) to generate the *apuA*::*spc* mutant. The triangles represent two sets of primers used for the detection of the disrupted *apuA* gene in recovered colonies indicating that homologous recombination with gDNA occurred after incubation in aPS, III) detection of recombinant mutant Δ*apuA* using *spc* (553 bp) and *apuA*::*spc* (614 bp) primers.

**Table 2:** Differential gene expression, shown as log2 fold-change values, of SS2 strain P1/*7* and SS9 strain 8067 grown in aPS compared to THY standard media.

| SS2 P1/7<br>Locus Tag | Gene | Functions | aPS vs THY |  |
| --- | --- | --- | --- | --- |
|  |  |  | SS2 P1/7 | SS9 8067 |
| SSU0049 | *comR* | Transcriptional activator of ComX | -2.42 | -0.73 |
| SSU0016 | *comX* | Master transcriptional regulator of the transformasome | -0.59 | -1.56 |
| SSU0061 | *cinA* | DNA binding and homologous recombination | -0.31 | 0.39 |
| SSU0062 | *recA* |  | -0.70 | -0.46 |
| SSU0924 | *radC* |  | -0.52 | -0.40 |
| SSU1083 | *coiA* |  | 0.66 | 0.36 |
| SSU0314 | *recG* |  | 2.49 | 1.34 |
| SSU1959 | *recF* |  | -2.56 | 1.21 |
| SSU0380 | *priA* |  | 0.86 | 0.69 |
| SSU0844 | *dprA* |  | 0.33 | 2.42 |
| SSU0126 | *comYA* | Transformasome apparatus | 4.02 | 0.47 |
| SSU0127 | *comYB* |  | 4.72 | 1.51 |
| SSU0128 | *comYC* |  | 5.48 | 4.36 |
| SSU0129 | *comYD* |  | 4.91 | 1.52 |
| SSU0130 | *comYE* |  | 3.05 | -0.48 |
| SSU0131 | *comYF* |  | 4.20 | 1.09 |
| SSU0132 | *comYG* |  | 4.26 | 0.47 |
| SSU0133 | *comYH* |  | -2.51 | 1.34 |
| SSU0610 | *comEA* | dsDNA receptor and channel | 2.07 | 0.33 |
| SSU0611 | *comEC* |  | 2.33 | 1.08 |
| SSU1009 | *endA* | competence associated endonuclease | -0.91 | -1.95 |
| SSU0144 | *ssbB* | ssDNA binding and protection | -1.34 | 1.14 |
| SSU1911 |  | fratricidin-like gene | 0.77 | 2.00 |
| SSU0039 |  | hypothetical protein | 1.97 | 0.04 |
| SSU0040 |  | CAAX amino protease | 1.15 | -0.08 |
| SSU0041 |  | membrane protein | 1.50 | 0.70 |
| SSU0042 |  | ABC transporter ATP-binding protein | 1.74 | 1.16 |
| SSU0043 |  | membrane protein | 1.60 | 1.58 |
| SSU0044 |  | ABC transporter ATP-binding protein | 1.91 | 1.84 |

SS strains cultured in human and porcine serum were efficiently transformed with plasmid or genomic DNA (gDNA) from 2 to 4 h, depending on initial bacterial density. Largest numbers of transformants were recovered in late exponential phase (Figure 1A, Table S1). Transformation efficiencies depended on SS serotype (i.e.: SS1, SS2, SS7, SS9 and SS14), presence of bacterial capsule and plasmid type (Table S1). In SS*7*, not transformable with its cognate XIP [9], natural competence could be induced by culturing in aPS (Table S1). High transformation efficiencies were obtained with zoonotic SS2 P1/*7* cultured in aPS and active human serum (aHS) but not when cultured in active Fetal Bovine Serum (aFBS) that contains lower levels of complement proteins and antibodies [16]. Serum heated for 30 minutes at 56°C to inactivate complement led to a remarkable reduction of competence suggesting that heat-labile serum components mediate efficient competence induction. Similar results were observed when (i) SS2 was cultured in serum depleted of serum components >10kDa by filtration; (ii) when sera had been stored for 2 weeks at 4°C and Fe^2+^ -chelating agents were added to aPS [12]; or (iii) when bacteria were grown in complex media not-supplemented with sugars [10] (mild starvation condition) (Table S1). To investigate if peptides mimicking XIP were present in serum we tested competence development with two SS2 mutants, Δ*comX* and Δ*comR*, which do not develop natural competence in the presence of XIP [9]. When cultured in aPS, transformation efficiencies of mutant and wild-type strains with pNZ8048 plasmid were similar, indicating that regulation of natural competence is independent of comR and comX. Transformation and homologous chromosomal recombination of a linear DNA fragment or gDNA from a Δ*apuA* mutant containing the *apuA*::*spc* gene was also verified by sequencing (Figure 1D, Table S1).

A mutant lacking pilus competence protein ComYC was either not transformable or yielded less than 5 transformants suggesting that the competence pilus is required for SS competence development in serum (Figure 1C, Table S1). To assess whether a similar mechanism exists in *S. pneumoniae* we grew strains A66 and 2127 in aHS, but obtained no transformants (unpublished data).

Here, we show that natural competence can be induced in potentially zoonotic SS2 strains when incubated cultured in aHS or aPS. Several SS strains and serotypes became naturally transformable with plasmid or gDNA in aPS including one strain that could not be transformed with the XIP allele encoded in its genome. The mechanism of competence development was ComR/ComX-independent and ComYC-dependent suggesting that serum-induced competence requires uptake of DNA by the transformasome.

Natural competence may allow uptake of foreign DNA at infection sites and contribute to bacterial survival and virulence [17]. Our preliminary results suggest SS exposure to complement factors may be involved in competence development. We have recently found in our lab that inflammatory stimuli induce, in intestinal organoids, expression of complement factors involved in early opsonisation and phagocyte chemotaxis [18], suggesting that bacteria colonising epithelia may also be exposed to complement factors. Novel strategies to inhibit natural competence may help to limit the spread of antibiotic resistance and pathogen evolution.

## Acknowledgments

The authors thank Edoardo Zaccaria for providing *com* mutant strains (*comR*, *comX* and *comYC*). This research was financially supported by EU Horizon 2020 Program Grant agreement ID 727966, funded under H2020-EU.3.2.1.1.

## Ethical standards

The authors have no conflict of interest to disclose.

