## Supplementary Table 1 for "Complement active human and porcine serum induces natural competence for genetic transformation in the emerging zoonotic pathogen *Streptococcus suis*"

**Transformation of SS in aPS and other media: CFU transformants per ml with 2 ug of plasmid or linear DNA**

**Transformation efficiencies of SS2 P1/7 grown in aPS at different bacterial density**

| Hours | serial dilution SS2 O/N culture in aPS | | |
|---|---|---|---|
| | 1/10 | 1/100 | 1/1000 |
| 1 H | 0 | 0 | 0 |
| 2 H | 165 | 310 | 0 |
| 3 H | 370 | 310 | 0 |
| 4 H | 185 | 400 | 340 |

**Transformation efficiencies of SS serotypes grown in aPS**

| Hours | SS serotypes | | | | |
|---|---|---|---|---|---|
| | SS1 strain 6388 | SS2 strain P1/7 | SS7 strain TMW_SS087 | SS9 strain 8067 | SS14 strain 13730 |
| 1 H | 125 | 0 | 0 | >1000 | >1000 |
| 2 H | 100 | 330 | 0 | >1000 | >1000 |
| 3 H | >1000 | 515 | 0 | 750 | 365 |
| 4 H | >1000 | 230 | 290 | 650 | 320 |

**Effect of *cps* on transformation efficiencies of SS2 strain S10 grown in aPS**

| Hours | | *cps gene mutant* |
|---|---|---|
| | SS2 s10 WT | SS2 s10 Δ*cps* |
| 1 H | 0 | 655 |
| 2 H | 330 | 605 |
| 3 H | 515 | 0 |
| 4 H | 230 | 0 |

**Effect of plasmid size in SS2 P1/7 on transformation efficiencies in aPS**

| Hours | Plasmid | |
|---|---|---|
| | pLABTarget_c | pNZ8048 |
| 1 H | 0 | >1000 |
| 2 H | 15 | 130 |
| 3 H | 370 | >1000 |
| 4 H | 110 | 0 |

**Transformation efficiencies of SS2 P1/7 grown in sera from different hosts**

| Hours | Host | | |
|---|---|---|---|
| | aPS | aHS | aFBS |
| 1 H | 0 | 0 | 0 |
| 2 H | 330 | 85 | 0 |
| 3 H | 515 | 280 | 0 |
| 4 H | 230 | 55 | 0 |

**Transformation efficiencies of SS2 P1/7 grown in PS inactivated and depleted of compounds >10kDa**

| Hours | Growth condition | | |
|---|---|---|---|
| | aPS control | iPS | aPS <10 kDa |
| 1 H | >1000 | 65 | 40 |
| 2 H | 130 | 80 | 5 |
| 3 H | 0 | 0 | 0 |
| 4 H | >1000 | 30 | 0 |

**Effect of storage period of aPS at 4°C on transformation efficiency of SS2 P1/7**

| Hours | 1 day @ 4°C | | 14 days @ 4°C | |
|---|---|---|---|---|
| | 1/10 | 1/100 | 1/10 | 1/100 |
| 1 H | 0 | 0 | 2 | 0 |
| 2 H | 165 | 310 | 15 | 0 |
| 3 H | 370 | 310 | 15 | 0 |
| 4 H | 110 | 400 | 65 | 0 |

**Transformation efficiencies of SS2 P1/7 grown in different media**

| Hours | Growth condition | | |
|---|---|---|---|
| | aPS control | THY | *Complex media |
| 1 H | **55** | <5 | 0 |
| 2 H | **1450** | <5 | 0 |
| 3 H | **0** | 0 | 0 |
| 4 H | **0** | 0 | 0 |

*** Complex media without sugars addition**

**Effect of the addtion or depletion of $Fe^{2+}$ on transformation efficiency of SS2 P1/7 grown in aPS and THY**

| Hours | aPS | | | THY (Negative control) | | |
|---|---|---|---|---|---|---|
| | control | + FeII | - FeII | control | + FeII | - FeII |
| 1 H | >1000 | 0 | <5 | <5 | 0 | 0 |
| 2 H | 130 | 35 | <5 | 0 | 0 | 0 |
| 3 H | 0 | 0 | 0 | 0 | 0 | 0 |
| 4 H | >1000 | >1000 | 0 | 0 | 0 | 0 |

**Transformation efficiencies of master regulators ComR/ComX deletion mutants and transposome ComYC mutants in SS2 strain S10 grown in aPS and THY**

| Hours | aPS | | | | THY (Negative control) | | | |
|---|---|---|---|---|---|---|---|---|
| | ΔcomX | ΔcomR | ΔcomYC | WT | ΔcomX | ΔcomR | ΔcomYC | WT |
| 1 H | 195 | 365 | <5 | 300 | 0 | 0 | 0 | <5 |
| 2 H | 365 | 250 | 0 | 50 | 0 | 0 | 0 | <5 |
| 3 H | 270 | 175 | 0 | 60 | 0 | 0 | 0 | 0 |
| 4 H | 45 | 75 | 0 | 85 | 0 | 0 | 0 | 0 |

**Transformation efficiencies of SS2 P1/7 with gDNA from an Δ*apuA* mutant strain**

| Hours | Recombination with DNA from *apuA::spc* | | | |
|---|---|---|---|---|
| | PCR | lysate 20 ul | lysate 50 ul | lysate 100 ul |
| 1 H | 150 | 0 | 170 | 240 |
| 2 H | 450 | 15 | 130 | 500 |
| 3 H | 685 | 0 | 210 | 230 |
| 4 H | 0 | 0 | 310 | 145 |
